# Coronavirus infection and PARP expression dysregulate the NAD Metabolome: an actionable component of innate immunity

**DOI:** 10.1101/2020.04.17.047480

**Authors:** Collin D. Heer, Daniel J. Sanderson, Lynden S. Voth, Yousef M.O. Alhammad, Mark S. Schmidt, Samuel A.J. Trammell, Stanley Perlman, Michael S. Cohen, Anthony R. Fehr, Charles Brenner

## Abstract

Poly-ADP-ribose polymerase (PARP) superfamily members covalently link either a single ADP-ribose (ADPR) or a chain of ADPR units to proteins using nicotinamide adenine dinucleotide (NAD) as the source of ADPR. While the well-known poly-ADP-ribosylating (PARylating) PARPs primarily function in the DNA damage response, many non-canonical mono-ADP-ribosylating (MARylating) PARPs are associated with cellular antiviral responses. We recently demonstrated robust upregulation of several PARPs following infection with Murine Hepatitis Virus (MHV), a model coronavirus. Here we show that SARS-CoV-2 infection strikingly upregulates MARylating PARPs and induces the expression of genes encoding enzymes for salvage NAD synthesis from nicotinamide (NAM) and nicotinamide riboside (NR), while downregulating other NAD biosynthetic pathways. We show that overexpression of PARP10 is sufficient to depress cellular NAD and that the activities of the transcriptionally induced enzymes PARP7, PARP10, PARP12 and PARP14 are limited by cellular NAD and can be enhanced by pharmacological activation of NAD synthesis. We further demonstrate that infection with MHV induces a severe attack on host cell NAD^+^ and NADP^+^. Finally, we show that NAMPT activation, NAM and NR dramatically decrease the replication of an MHV virus that is sensitive to PARP activity. These data suggest that the antiviral activities of noncanonical PARP isozyme activities are limited by the availability of NAD, and that nutritional and pharmacological interventions to enhance NAD levels may boost innate immunity to coronaviruses.

---

Disease attributed to the current novel coronavirus (CoV) outbreak (COVID-19) has rapidly spread globally, infecting more than 33 million people and killing more than a million as of late September, 2020 (1). The causative agent, severe acquired respiratory syndrome coronavirus 2, SARS-CoV-2, is transmitted largely by aerosol and liquid droplets that infect cells of the lung epithelium (2). Severe disease is thought to proceed through a combination of robust viral replication and a cytokine storm in which host inflammation damages multiple organ systems. While many therapeutic approaches are under investigation, the evidence-basis for effective prevention and treatment agents remains limited.

CoV genomes do not encode enzymes needed for ATP generation, nucleotide, amino acid, lipid or protein synthesis, and therefore depend on exploitation of host functions to synthesize and assemble virus (3-5). Cellular and viral energy generation and biosynthetic programs depend on the four nicotinamide adenine dinucleotide (NAD) coenzymes, NAD^+^, NADH, NADP^+^ and NADPH, which are the central catalysts of metabolism (6). These coenzymes accept and donate electrons in essential, ubiquitous processes of fuel oxidation, lipid, nucleotide and amino acid biosynthesis, and the generation and detoxification of reactive oxygen species. The specific roles of these coenzymes in viral replication and antiviral defenses are largely unexplored.

Several members of the PARP superfamily are interferon-stimulated genes (ISGs) that have been implicated in restriction of viral replication through mechanisms that are not well understood (7,8). With the exception of the enzymatically inactive PARP13 protein, PARP isozymes have an absolute requirement for NAD^+^ (9,10). However, rather than using NAD^+^ as an electron acceptor, PARPs use NAD^+^ as an ADPR donor in protein modification reactions (6). The canonical and best characterized PARP isozymes, PARP1 and PARP2, form PAR largely in response to DNA damage. However, most other members of the PARP superfamily exhibit MARylation activities on target proteins (9). We previously showed that MHV infection strongly induces expression of noncanonical PARP isozymes PARP7, PARP9, PARP10, PARP11, PARP12, PARP13 and PARP14 (11). To determine whether these gene expression changes are incidental to MHV infection, facilitate MHV infection, or are part of an innate immune response against MHV, we treated cells with siRNAs to knock down expression of these genes and then analyzed the impact on MHV replication. Our data showed that PARP7 plays a role in facilitating replication. In contrast, PARP14 is required for full induction of interferon (IFN)-β expression (11,12), suggesting that PARP14 is directly involved in establishing the innate immune response in CoV-infected cells.

Most CoV genomes encode 16 non-structural proteins (nsps) (3-5). Nsp3 contains a macrodomain, herein termed the CoV ADPR hydrolase (CARH) that removes ADPR modifications from acidic amino acids on protein targets. Thus, CARH reverses the modification that is installed by the IFN-induced activities of MARylating PARP isozymes (13). CARH activity is required for virulence *in vivo* using mouse models of both MHV and SARS-CoV (13-15). Moreover, an active site mutation that ablates CARH activity in MHV resulted in a virus that replicates poorly in primary bone-marrow derived macrophages (BMDMs) (11). We further identified PARP12 and PARP14 as CoV-induced ISGs that are required for depressed replication of CARH mutant viruses, indicating that their activity is opposed by CARH-mediated reversal of MARylation (11).

In support of the antiviral roles of MARylating PARP isozymes, PARP12 was shown to promote degradation of nsp1 and nsp3 in Zika virus infection (16). PARP12 has also been shown to inhibit a wide variety of RNA viruses, including several alphaviruses, which also contain nsp3-encoded ADPR hydrolase activities (17,18). These observations suggest that key events in the innate immune response to viral infections are played out in the infected cell’s NAD metabolome.

To determine whether the dramatic upregulation of PARPs seen following MHV infection is conserved among CoVs and also whether cellular infection by CoVs disturbs NAD homeostasis, we analyzed transcriptomic data from SARS-CoV-2-infected human cell lines and organoids, SARS-CoV-2-infected ferrets, a lung biopsy from people living with or dead from COVID-19, and bronchoalveolar lavage fluid (BALF) from healthy individuals as well as those infected with COVID-19. These data indicate that the same noncanonical PARPs induced by MHV are induced by SARS-CoV-2 and that *in vivo* infection with SARS-CoV-2 down-regulates synthesis of NAD from tryptophan and nicotinic acid (NA) while up-regulating synthesis capacity from NAM and NR. We further show that disturbances to the NAD transcriptome scale with viral load. Though noncanonical PARP isozymes are known to use NAD^+^ to MARylate target proteins, it had not been reported that expression of these enzymes can drive down cellular NAD^+^. Here we show that PARP10 overexpression is sufficient to depress cellular NAD^+^ and that the MARylating activities of PARP10, PARP12 and PARP14 can be pharmacologically increased by enhancing NAD salvage synthesis with SBI-797812 (SBI), a NAMPT activator (19).

Though the essentiality of CARH for viral pathogenesis argues for cellular NAD and noncanonical PARP induction as antiviral, it remained conceivable that a depressed cellular NAD metabolome is an adaptive antiviral response to restrict viral biosynthetic processes. We therefore established a cellular system to test whether increased NAD status opposes MHV infection. Consistent with our transcriptomic analysis that CoV infection downregulates NA salvage and upregulates NAM and NR salvage, we found, using an MHV mutant virus sensitive to PARP activity, that NA minimally inhibited viral replication, while NAM, SBI and a clinically tested preparation of NR (Niagen)(20-22) strongly inhibit MHV replication. The data justify further analysis of how nutritional and therapeutic modulation of NAD status may potentially restrict viral infection by boosting PARP activity.

## RESULTS

### SARS-CoV-2 Infection of Human Lung Systems Induces a Noncanonical PARP Isozyme Transcriptional Program

MHV infection in murine BMDMs launches a transcriptional program that induces transcription of noncanonical PARP isozymes PARP7, PARP9, PARP10, PARP11, PARP12, PARP13 and PARP14 by > 5-fold (11,12). To determine whether SARS-CoV-2 also dysregulates the NAD system upon infection, we assembled and analyzed a set of 71 genes that encode the enzymes responsible for conversion of tryptophan, NA, NAM, and NR to NAD^+^, plus the enzymes responsible for NAD(H) phosphorylation, NADP(H) dephosphorylation, NAD^+^-dependent deacylation, MARylation, PARylation, cADP-ribose formation, nicotinamide methylation/oxidation, and other related functions in transport, binding, redox and regulation (**Supplementary Material 1**). We then analyzed RNAseq data from SARS-CoV-2 infection of three human lung cell lines, Calu-3, normal human bronchial epithelia (NHBE), and A549 (**Fig. 1A-C**) (23). Calu-3 cells exhibit the most robust response, inducing PARP7, PARP9, PARP10, PARP12 and PARP14 more than 4-fold as well as PARP3, PARP4, PARP5a, PARP5b and PARP8 to a lesser but statistically significant degree (**Fig. 1A**). In NHBE cells, SARS-CoV-2 induces transcription of PARP9, PARP12 and PARP14, while in A549 cells it induces PARP9, PARP10, PARP12 and PARP14, with lesser effects on PARP7 and PARP13 (**Fig. 1B-C**).

**Figure 1.**
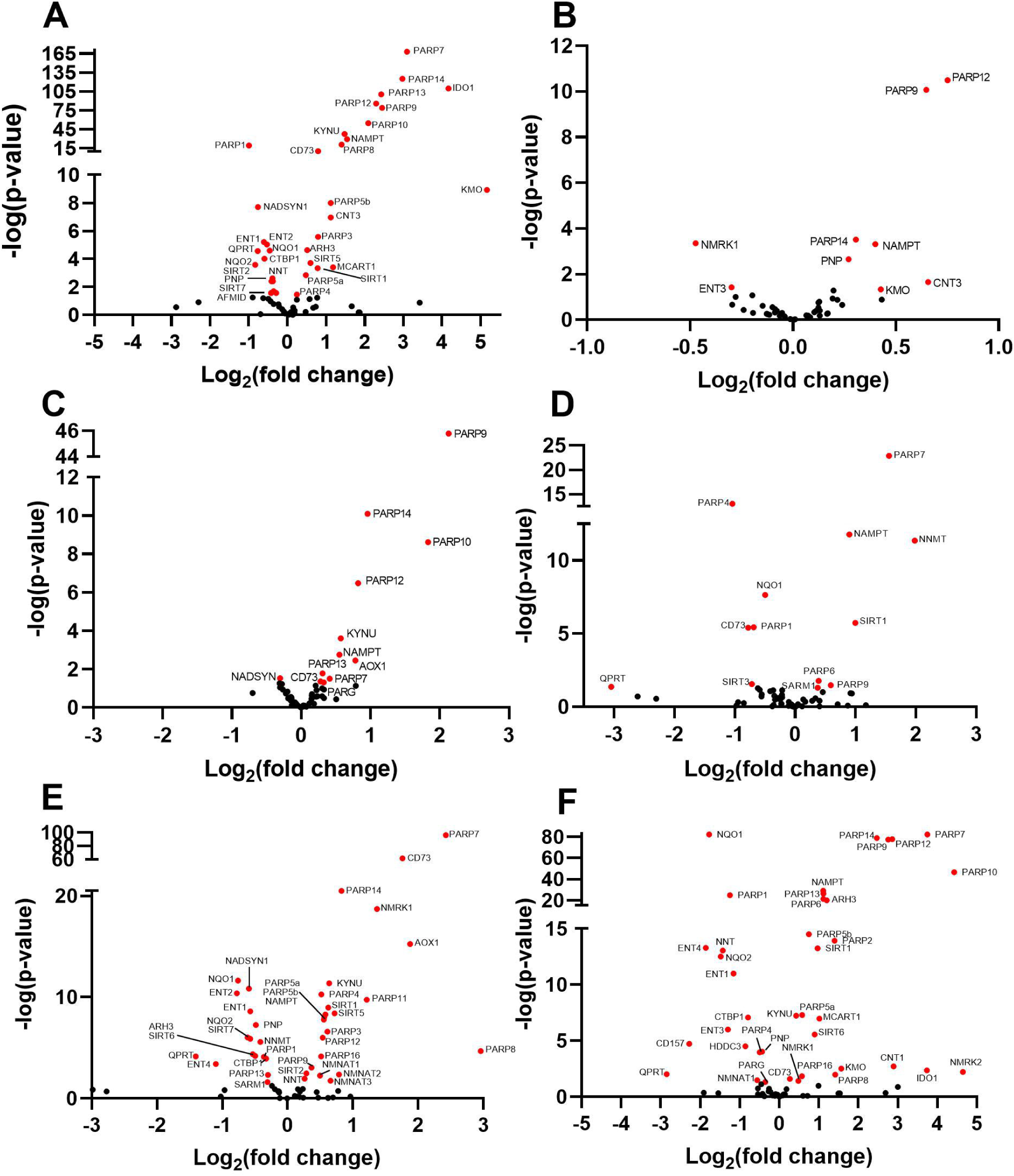
SARS-CoV-2 dysregulates the NAD gene set *in vitro* as a function of viral load. Differential expression analysis was performed on RNAseq data with respect to a 71 gene set representing the NAD transcriptome (Supplementary Table 1). Depicted are Volcano Plots representing normalized relative expression versus -log(P) with respect to mock infected in A) human Calu3 lung cancer cells (MOI = 2), B) NHBE cells (MOI = 2), C and D), A549 cells at low MOI without and with introduction of ACE2 expression, respectively, E and F) A549 cells at high MOI = 2 without and with introduction of ACE2 expression. Further information is available in Supplementary Materials 2-7.

### Transcriptional Dysregulation of NAD Metabolism Scales with Viral Titer

Angiotensin-converting enzyme 2 (ACE2) serves as a receptor for SARS-CoV-2 entry (24). To determine the effect of overexpression of ACE2 or greater viral exposure on the NAD transcriptome, we compared high quality RNAseq data from control or ACE2-overexpressing A549 cells infected at low (0.2) or high (2) multiplicity of infection (MOI). Greater viral exposure dysregulated more NAD related genes (**Fig. 1D-F)**. The viral load-dependent changes came in three types. First, PARP7 is strongly dependent on viral load, being minimally induced in A549 cells infected at a low MOI (**Fig 1C**). However, in cells expressing ACE2 or infected at high MOI, PARP7 was one of the most highly induced PARP isozymes (**Fig. 1D-F**). Second, as ACE2 or more virus was added, transcription of more PARP isozymes was disturbed. ACE2 and higher MOI both downregulated expression of PARP1 while they upregulated expression of 12 of the other 16 PARP superfamily members (**Fig. 1F**). Third, on the basis of transcriptional changes, viral infection alters expression of NAD biosynthesis pathways. ACE2 and higher MOI both downregulated expression of QPRT, which is required for the conversion of quinolinate to nicotinic acid mononucleotide in the *de novo* biosynthetic pathway that originates with tryptophan. Similarly, NADSYN, which is required for synthesis of NAD from both tryptophan and NA is down-regulated by viral infection, suggesting that CoV infection might disadvantage repletion of the NAD metabolome from either tryptophan or NA. In contrast, NAMPT, which is required for NAD synthesis from NAM, was consistently upregulated in SARS-CoV-2 infected cells, and NMRK1, which is required for NAD synthesis from NR, was up-regulated when cells were infected at high MOI. In addition, the normally minimally expressed NR kinase gene, NMRK2, was upregulated in the ACE2 overexpressing cells infected at high MOI.

**Table 1.** Impact of NAD-boosting compounds on N1347A replication. The average fold reductions in viral titer of N1347A in 2 independent experiments on 17Cl-1 or BMDM cells. The p values are from an unpaired two-tailed t-test of data from both experiments combined.

| Compound | Fold Reduction in N1347A Viral Titer | p value |
| --- | --- | --- |
| NA | 1.9 | 4.4E-2 |
| SBI | 4.4 | 1.7E-3 |
| NAM | 8.3 | 6.3E-4 |
| NR | 6.4 | 8E-5 |
| NR (BMDM) | 2.7 | <1E-6 |

### Primary 3-D cell culture and in vivo SARS-CoV-2 Infection Disturb Noncanonical PARP Isozyme Expression and NAD Biosynthesis

Based on the finding that SARS-CoV-2 infects gut enterocytes, we analyzed RNAseq data obtained from infected enterocyte organoids (25). Similar to the lung cell lines and to the transcriptional effect of MHV infection, enterocyte organoids infected with SARS-CoV-2 induce PARP9, PARP12 and PARP14 (**Fig. 2A**).

**Figure 2.**
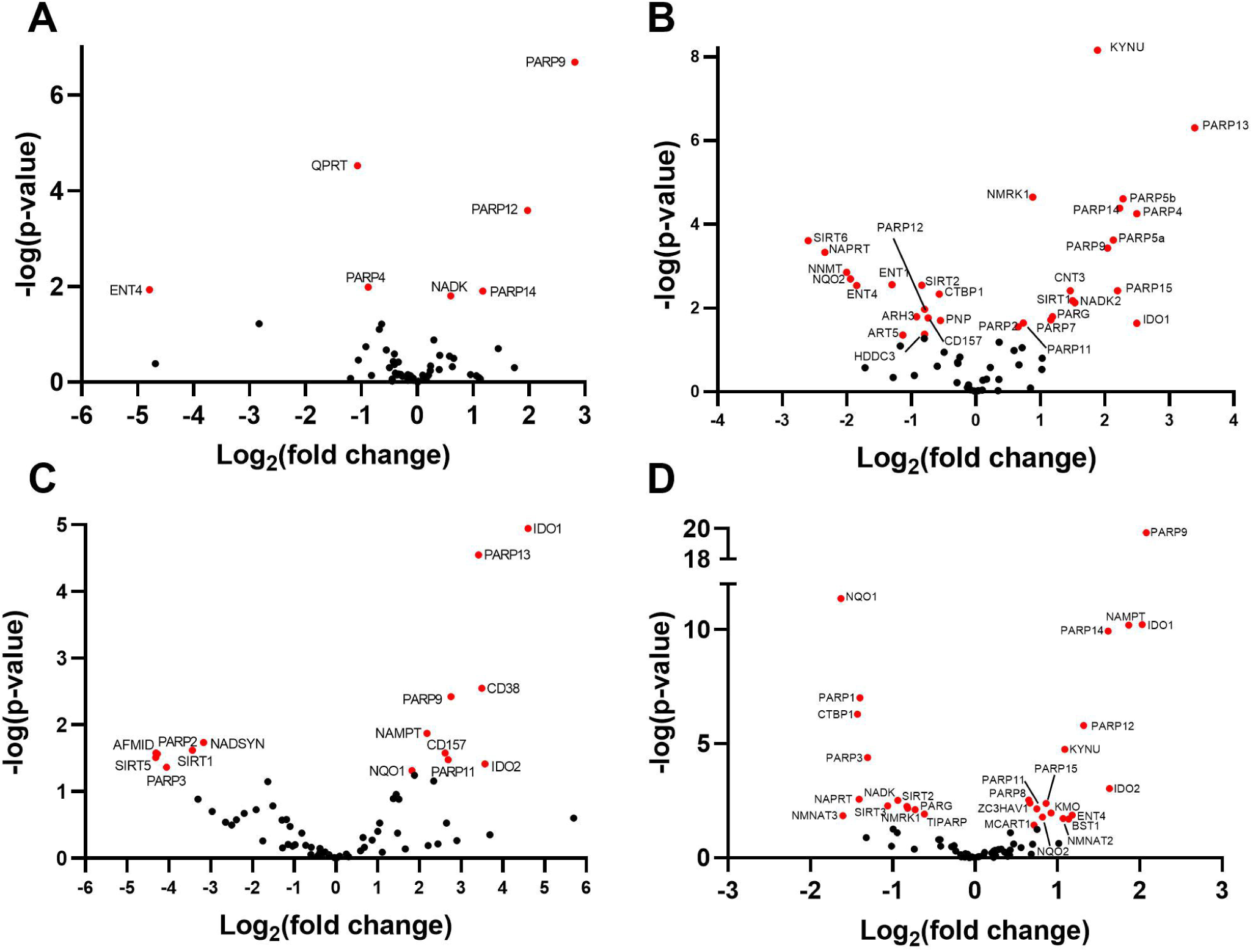
SARS-CoV-2 dysregulates the NAD gene set *in vivo*. Differential expression analysis was performed on RNAseq data with respect to mock infected in A) expanding enterocytes (MOI=1), B) ferret trachea infected with SARS-CoV-2, C) lung of a diseased COVID-19 patient versus a control lung sample and D) BALF from SARS-CoV2 infected versus healthy control human patients. Further information is available in Supplementary Materials 8-11.

Ferrets have been shown to be permissive to SARS-CoV-2 infection (26) and are being used as a system to probe host responses as well as potential preventative and therapeutic agents. To determine if PARP upregulation following SARS-CoV-2 infection is also observed *in vivo*, we probed high-quality RNAseq data from the tracheas of mock and 3-day SARS-CoV-2 infected ferrets (23) and found that the noncanonical PARP induction program is conserved in this relevant animal model (**Fig. 2B**). Specifically, PARP4, PARP5, PARP9, PARP13, PARP14 and PARP15 were all > 4-fold induced with significant but lesser induction of PARP7 and PARP11. In ferret tracheas, we observed other significant alterations to the NAD gene set. The data indicate that transcription of NMRK1 and concentrative nucleoside transporter CNT3 are induced in response to CoV infection, suggesting increased capacity for uptake and conversion of NR to NAD^+^ and NADP^+^ (27). Notably, in mouse models of damaged brain and heart, upregulation of NMRK gene expression is associated with therapeutic efficacy of NR (28,29). Additionally, the ferret data show strongly depressed NNMT expression—by decreasing NAM methylation, this gene expression change could promote NAM salvage (30) and the efficiency of NAD-boosting by NR or NAMPT activators (20), representing a homeostatic attempt of virally infected cells to maintain their NAD metabolome (31).

### COVID-19 Patient Samples Recapitulate the PARP Induction Program Seen in Ferrets and In Vitro

Finally, though ferrets are susceptible to infection by SARS-CoV-2, they do not progress to the serious disease seen in humans (26). We therefore examined the NAD gene set in RNAseq data from the lung of an individual who died of COVID-19 (23). Though lacking the replicates and the synchrony of the ferret RNAseq data, the human data showed that PARP9, PARP11 and PARP13 were upregulated in the deceased individual’s lung (**Fig. 2C**).

Finally, we analyzed RNA-seq data obtained from the BALF of 430 SARS-CoV-2 infected people and 54 controls (32). Consistent with previous findings, PARP9 was upregulated nearly 4-fold while PARP12 and PARP14 were upregulated more that 2-fold (**Fig. 2D**).

Notably, NAD biosynthetic gene changes were conserved *in vivo* with NAMPT and NMRK1 gene expression increased by viral infection in ferrets, and NAMPT increased in the human patient samples. Further data on transcriptomic alterations to NAD synthesis in all nine data sets are provided in **Supplementary Material 12**.

### PARP10 Overexpression is Sufficient to Depress NAD^+^ Levels

It is well known that PARP1 activation by DNA damage greatly increases its catalytic activity, leading to depression of cellular NAD^+^ and ATP (9,10). It is less clear whether transcriptional induction of the MARylating enzymes such as PARP10 that are induced substantially by viruses including SARS-CoV-2 might disturb cellular NAD^+^. To test whether overexpression of a MARylating enzyme is sufficient to disturb the NAD metabolome, we performed targeted quantitative NAD analysis with LC-MS/MS using internal ^13^C standards (33) on HEK 293T cells expressing either GFP or a GFP-PARP10 fusion. We found that overexpression of GFP-PARP10 significantly depressed NAD^+^ compared to overexpression of GFP alone (**Fig. 3A**). We next determined if the PARP10-mediated loss in NAD^+^ could be restored by increasing cytosolic NAD^+^ synthesis with SBI, a small molecule activator of NAMPT, which promotes NAM salvage (19). SBI increased steady-state levels of NAD^+^ in GFP-expressing cells, but did not significantly boost NAD^+^ in PARP10-expressing cells, indicating that PARP10 expression is sufficient to limit cellular NAD^+^ (**Fig. 3A**).

**Figure 3.**
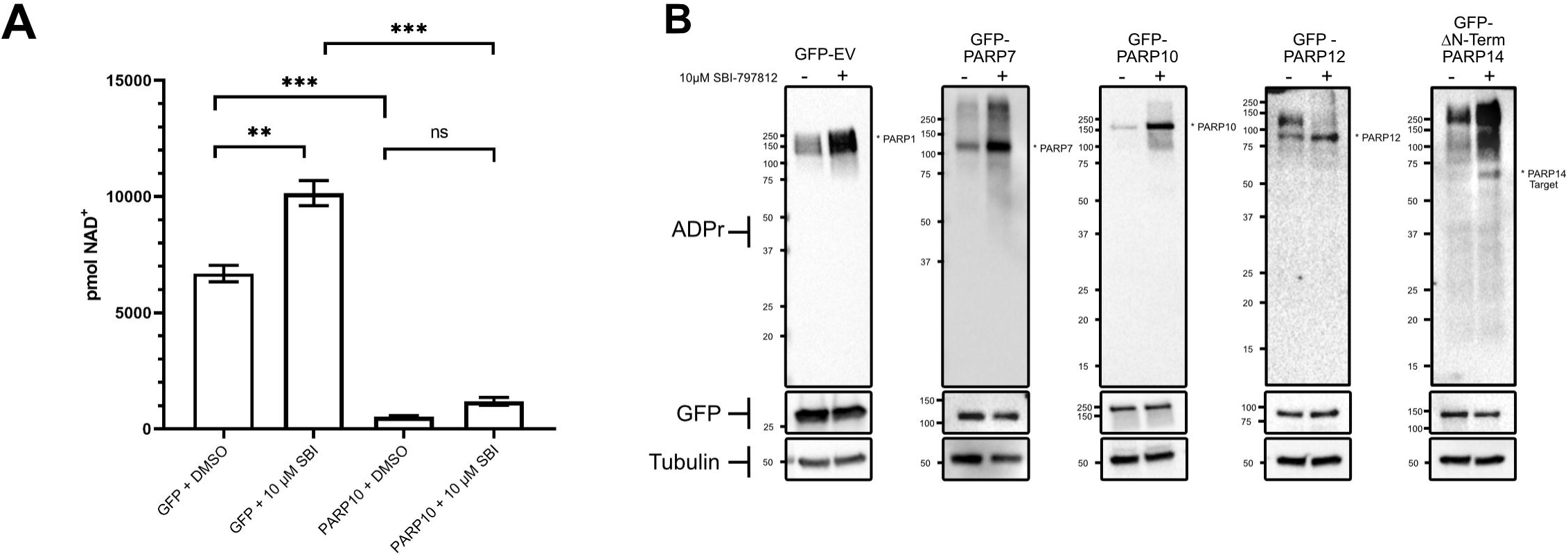
PARP10 overexpression is sufficient to depress cellular NAD^+^ levels while SBI enhances activities of overexpressed PARP7, PARP10, PARP12 and PARP14. A) HEK 293T cells were grown with the indicated expression plasmids for GFP or PARP10 and treated with NAMPT activator (10 μM SBI). n = 3 for each group. Error bars represent SEM, p-values are from an unpaired two-tailed t-test. See also Supplementary Information 12. B) GFP, PARP7, PARP10, PARP12 and PARP14-expressing HEK293 cells were treated with 10 μM SBI and cells were collected 18 hours later. Western blot using indicated antibodies indicate that SBI promotes PARP7, PARP10, PARP12 and PARP14 activity. n=3. Representative blots of three independent experiments are shown. ** p≤ 0.01, *** p≤ 0.001

### Enhanced NAD Salvage Increases the Activity of PARP Isozymes Induced by MHV & SARS-CoV-2 infection

The ability of SBI to elevate cellular NAD^+^ in GFP-expressing cells but not in PARP10-expressing cells suggested that overexpressed PARP isozymes are active at lower levels of cellular NAD^+^ but left open the question of whether overexpressed PARP isozymes have enzymatic activities that are limited by the depressed NAD^+^ status that they confer. To determine whether the activity of PARP7, PARP10, PARP12 and PARP14 can be increased by NAMPT stimulation, we measured ADPR-modified protein levels by western blot in HEK 293T cells overexpressing GFP, or GFP-tagged PARP7, PARP10, PARP12 or PARP14 in the presence or absence of SBI. Consistent with the ability of SBI treatment to elevate cellular NAD^+^, SBI enhanced PARP1 autoPARylating activity in GFP-expressing cells. SBI treatment resulted in a striking increase in PARP10 activity as evidenced by enhanced PARP10 autoMARylation. A similar result was observed with PARP7 and PARP12 albeit to a lesser degree. Finally, SBI treatment of PARP14-expressing cells resulted in a significant increase in PARP14 target MARylation (auto-MARylation was not detected under these conditions) (**Fig. 3B**). These results indicate that activities of overexpressed PARP isozymes are limited by cellular NAD^+^ levels and can be enhanced pharmacologically.

### MHV Infection Drives Down Cellular NAD^+^ and NADP^+^ in Infected Cells

Given that CoVs consistently induce members of the PARP superfamily at the mRNA level, we asked whether MHV infection alters the NAD metabolome. MHV is a model CoV that can be propagated under BSL-2 conditions, allowing us to obtain samples needed for quantitative analysis of the NAD metabolome. We infected delayed brain tumor (DBT) cells with MHV-A59 at a MOI of 3 and subjected the cells to quantitative targeted NAD metabolomics 12 hours after infection (33). Infection led to a > 3-fold depression of cellular NAD^+^ and NADP^+^ with respect to control after a mock infection (**Fig. 4A**). To address the possibility that these effects are specific to a cancer cell line and not a primary viral target cell population, we prepared bone marrow derived macrophages (BMDM) from C57BL/6 mice and infected them with MHV-A59 at a MOI of 3. Similar to DBT cells, we observed a greater than 3-fold depression of cellular NAD^+^ and NADP^+^ with respect to control cells at 12 hours (**Fig. 4B**). Thus, in primary cells that represent an authentic CoV target cell population, MHV depresses levels of NAD coenzymes.

**Figure 4.**
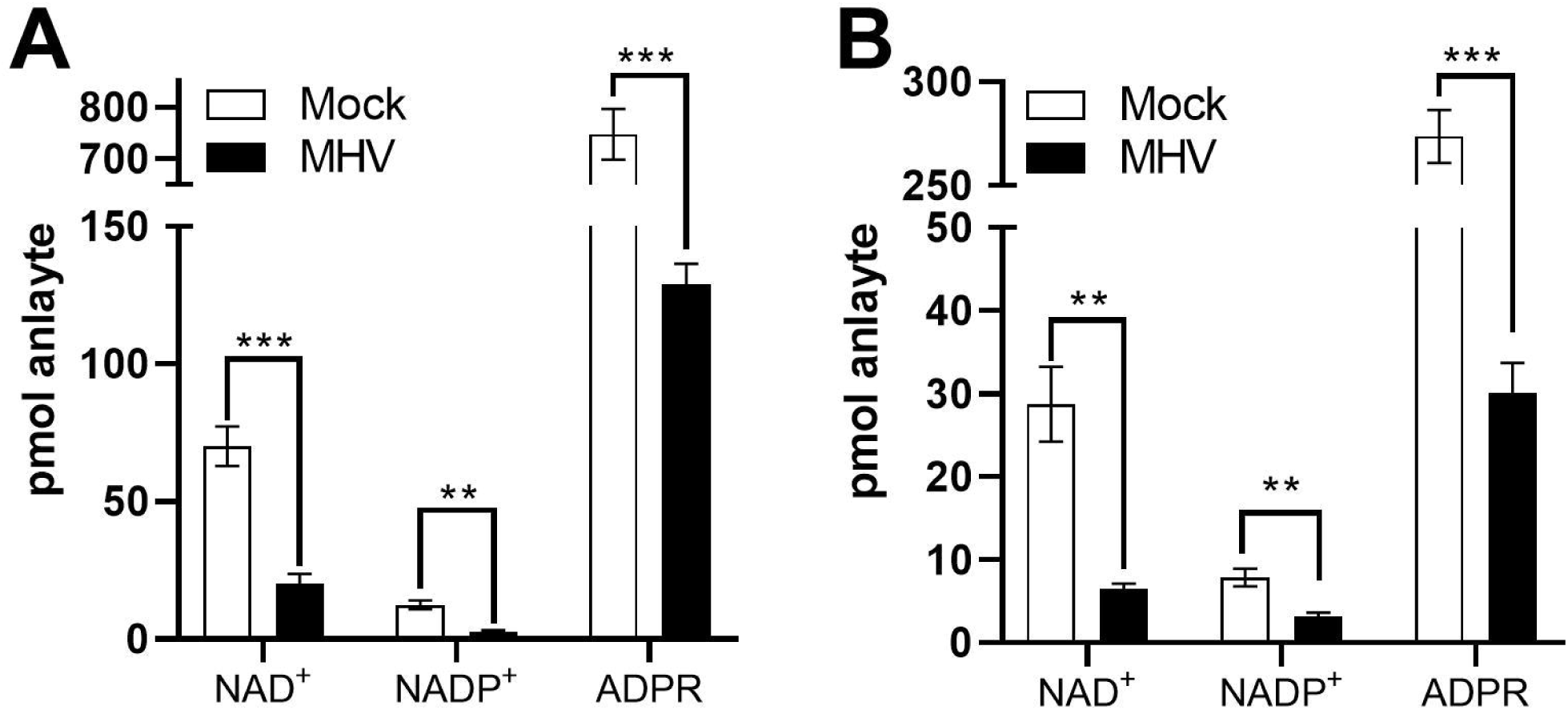
MHV infection disturbs the NAD metabolome. A) DBT cells and B) BMDM cells were mock infected or infected with MHV at a MOI of 3 PFU/cell and cells were collected at 12 hrs post-infection. n = 3-4 Mock; n = 4 MHV. Error bars represent SEM, p-values are from unpaired two-tailed t-test. **p ≤ 0.01, ***p ≤0 .001. See also Supplementary Materials 13-14.

### NAD Boosting Compounds Decrease Replication of a CARH-mutant MHV

We previously reported that the CARH domain of MHV and SARS-CoV Nsp3 proteins is required for maximum replication and pathogenesis *in vivo* (13-15). Moreover, the N1347A active site mutation that ablates the ADPR hydrolase activity of CARH and *in vivo* infectivity of MHV also resulted in a virus that replicates poorly in primary BMDM cells but reaches similar peak titers as WT virus in a transformed 17Cl-1 fibroblast cell line (11,14). These data suggest that a CARH inhibitor would be completely antiviral *in vivo* and would show some activity in dampening replication in cellular infection models.

Our data established that noncanonical PARPs are consistently induced by CoV infection (**Figs. 1-2**); can depress cellular NAD^+^ and have their MARylation activities limited by cellular NAD^+^ (**Fig. 3**); and have known antiviral activities (16-18). These data suggest that boosting cellular NAD through the transcriptionally upregulated NAMPT and NMRK pathways would have antiviral activity. However, it remained a possibility that the anti-viral response of infected cells evolved to drive down cellular NAD^+^ and NADP^+^ in order to rob invading viruses of biosynthetic capacity.

To test the hypothesis that NAD-boosting interventions would depress viral replication in sensitive cellular assays, we measured MHV N1347A infection of BMDM and 17Cl-1 systems as a function of addition of NA, NAM, SBI or NR (Niagen) (**Fig. 5A**). As previously reported, N1347A reached similar peak titers as WT virus in 17Cl-1 cells. However, the addition of NA, NAM, SBI and NR all significantly decreased its replication (**Fig. 5B**). Consistent with increased NAMPT and NMRK gene expression and depressed NADSYN expression in SARS-CoV-2 -infected cells (**Fig. 1-2**), NAM, SBI and NR had significantly greater effects on N1347A replication than NA. NAM, SBI and NR decreased N1347A replication by 8.3, 4.4, and 6.4 fold respectively, while NA decreased its replication by only 1.9 fold (**Fig. 5B, Table 1**). Consistent with the permissivity of 17Cl-1 cells to infection (11,14), these treatments did not affect WT virus replication in 17Cl-1 cells.

**Figure 5.**
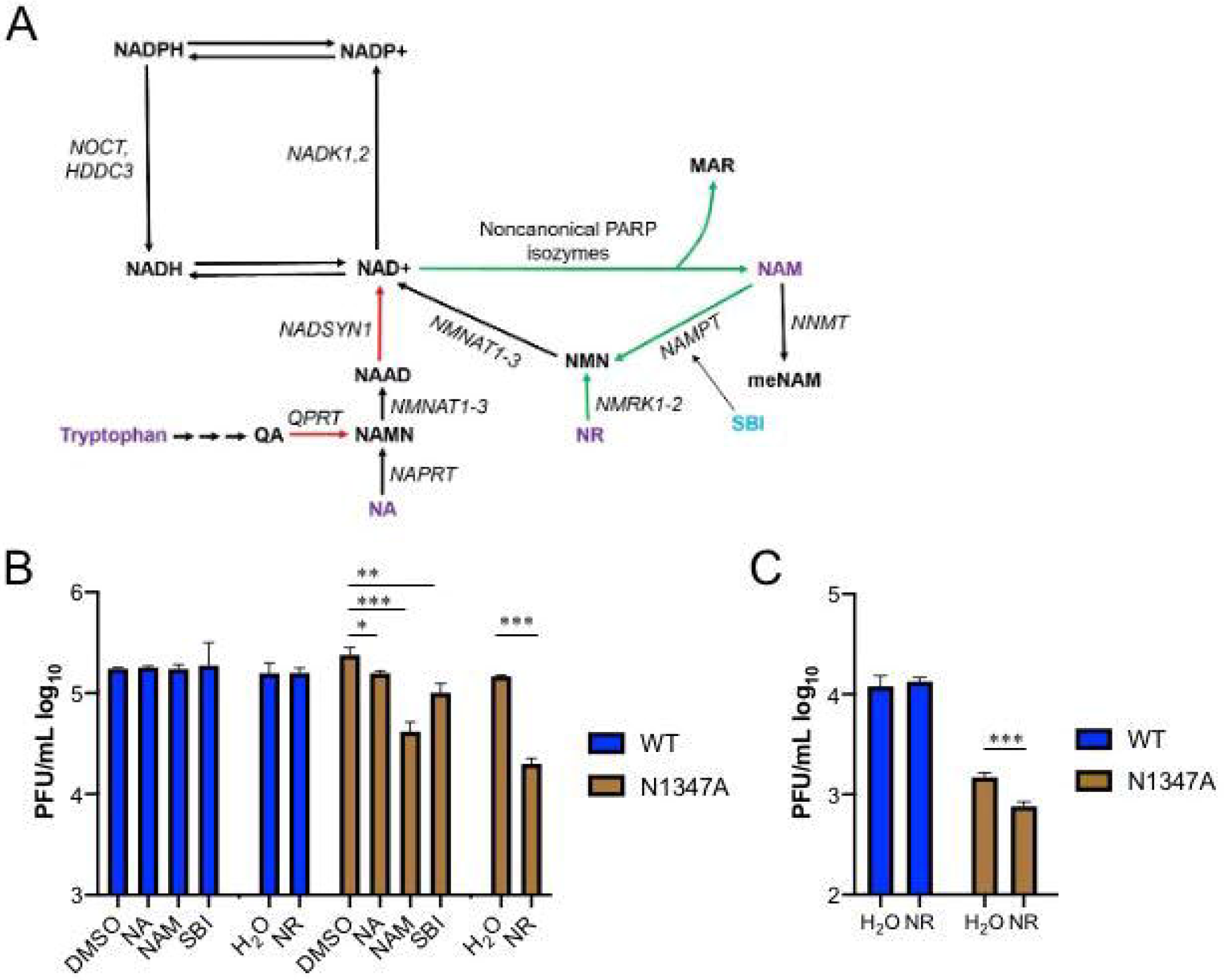
Boosting NAD^+^ levels depresses replication of CARH mutant MHV. A) NAD biosynthetic pathways. Red arrows depict gene expression that is depressed by SARS-CoV-2. Green arrows depict gene expression that is increased by SARS-CoVo2. B) 17Cl-1 cells were infected with 0.1 PFU/cell WT or N1347A MHV and either mock treated (DMSO or H2O) or treated with NA, NAM, SBI or NR as described in Methods. DMSO served as a solvent control for NAM, SBI and NA, while H_2_O served as a solvent control for NR. Cells were collected at 18 hpi and analyzed for virus replication by plaque assay. Data are representative of two independent experiments, n = 3 biological replicates. C) BMDMs were infected with 0.1 PFU/cell WT or N1347A MHV and treated with H_2_O or treated with NR as described in Methods. Cells were collected at 18 hpi and analyzed for virus replication by plaque assay. Data are representative of two independent experiments. n = 4 biological replicates. *p ≤ 0.05, **p ≤ 0.01, ***p≤0.001.

Next we tested the ability of NR to reduce N1347A replication in BMDM cells. Here, N1347A with no treatments had a replication defect of 4.8-fold compared to WT virus, similar to our previous report (11). Consistent with the view that higher NAD^+^ status depresses viral infectivity in cellular assays, addition of NR to cultures infected by N1347A further depressed replication by 2.7-fold (**Fig. 5C, Table 1**).

Our data indicate that: *i)* CoV infection initiates the expression of multiple noncanonical PARP isozymes and dysregulates other genes involved in NAD metabolism; *ii)* CoV infection and PARP10 expression can dramatically decrease NAD^+^ accumulation; and *iii)* NAD^+^ boosting agents can improve PARP isozyme function and decrease the replication of a CoV that is sensitive to MARylating activities.

## DISCUSSION

SARS-CoV-2 is a highly infectious agent that constitutes a severe threat to public health (2). Morbidity and mortality data make it clear that age, smoking status and multiple preexisting conditions greatly increase the frequency of serious illness and death (34). There is an abundance of data from model systems and humans that age and conditions of metabolic stress including obesity and type 2 diabetes (35), smoking (36), heart failure (29), nerve damage (37) and central brain injury (28) challenge the NAD system in affected tissues. Though PARP1 was known to be a significant consumer of NAD^+^, we showed that noncanonical PARP isozymes are consistently upregulated by CoV infections, that PARP10 overexpression can depress the NAD metabolome, and that four noncanonical PARP isozymes that are induced by SARS-CoV-2 have MARylating activities that are limited by cellular NAD status. In addition, the NAD metabolome was depressed by MHV infection. While the degree of depression of the NAD metabolome is surely sensitive to time and MOI, the >3-fold depression seen in this study is precedented in human immunodeficiency virus and Herpes virus infections (38,39).

Though the genetic requirement for CARH for viral replication *in vivo* (11,13-15) strongly suggested that higher NAD^+^ status would be protective, it was also conceivable that PARP-mediated cellular repression of the NAD metabolome constituted a mechanism for cells to deprive the virus of anabolic capacity. However, here we showed that boosting NAD^+^ through the NR and NAMPT pathways depresses replication in a CARH-mutant cellular infection model with no effect on replication of wild-type virus. Further experiments will be required to identify the key MARylated targets that are potentially enhanced by NAD-boosting and reversed by CARH activities.

Based on gene expression data in response to SARS-CoV-2 infection (**Fig. 1-2**), NAD boosting approaches involving increased *de novo* or NA-dependent synthesis are unlikely to be strongly effective because they require expression of genes such as QPRT, NADSYN and NAPRT (40) that are depressed by SARS-CoV-2 infection. Consistent with gene expression changes, in the cellular infection system, we showed that addition of NA only modestly reduced virus replication (**Fig. 5B-C, Table 1**).

Based on gene expression data as well as MHV cellular infection data, NAM, NAMPT activators and NR have similar potential to be strongly protective *in vivo*. However, there are a few caveats. First, that at pharmacological doses, NAM has the potential to function as a PARP inhibitor (41). Second, NAMPT is considered a driver of pulmonary vascular remodeling and potentially a target to be inhibited to maintain lung health of some people at risk for COVID-19 (42). Thus, in order to maximize the likelihood of success in human CoV prevention and treatment trials, care should be taken to carefully compare efficacy and dose-dependence of NR, SBI and NAM with respect to control of cytokine storm and antiviral activities *in vivo*.

The cellular results presented herein warrant the testing of NAD boosting agents in the context of *in vivo* CoV infections. In addition to animal trials, the safety of various forms of vitamin B3 should allow rapid clinical assessments of NAD boosters to be evaluated in two placebo-controlled contexts.

First, we suggest that improved NAD status could help blunt the severity of infection by sustaining PARP-dependent IFN signaling in the face of the self-limiting nature of cellular NAD during infection and by limiting the storm of inflammatory cytokines that is typically associated with serious disease (43). In a small placebo-controlled clinical trial designed to address the oral safety and activity of Niagen NR in older men, it was discovered that 1 gram of NR per day depresses levels of IL-6, IL-5, and IL-2 (22). Based on these findings, we suggest that NAD boosters be tested on hospitalized and nonhospitalized COVID-19 patients with primary endpoints of disease recovery, oxygenation and cytokine levels. Similar approaches might also be tested in the context of other viral infections that induce noncanonical PARP isozymes and/or encode viral ADP-ribosylhydrolase activities (17,18).

Second, as there is no standard of care for housemates or caretakers of infected people, we suggest that NAD-boosting could be tested as a placebo-controlled intervention for people in the proximity of quarantined or hospitalized COVID-19 individuals—the primary endpoint would be protection against infection.

While caution should be exercised with respect to any preventative measure, NAD boosting approaches have the potential to support the innate immune system and address the age-, smoking- and comorbid conditions associated with worse SARS-CoV-2 outcomes (34). The potential societal benefit of a safe and readily available molecule to support prevention and public health is hard to overstate, especially as new outbreaks of COVID-19 emerge.

## EXPERIMENTAL PROCEDURES

### RNASeq analysis

Sequence counts for **Fig. 1** including cell lines infected with SARS-CoV-2 (strain USA-WA1/2020) were derived from published RNAseq (GSE147507) (23). Briefly, A549 (+/-ACE2), Calu3 and NHBE cell lines were infected with either Mock or SARS-CoV-2 virus for 24 hours. Data for **Fig. 2** were drawn from several sources. First, expanding enterocyte data (**Fig. 2A**) were obtained from Supplementary Table 2 of (25). Briefly, organoids were infected with SARS-CoV-2 at an MOI of 1 in expansion medium for 60 hr. RNA was collected, and differential gene expression analysis was performed using the DESeq2 package. The ferret (**Fig. 2B**) and deceased human data (**Fig. 2C**) were also obtained from GSE147507. Ferrets were infected with SARS-CoV-2 at a PFU of 5 x 10^4^. More information regarding sample preparation and data processing can be found at GEO accession GSE147507. Finally, the human BALF data **Fig. 2D** were obtained from GSE152075. Briefly, samples were obtained from nasopharyngeal swabs of from 430 male and female individuals with SARS-CoV-2 infection and 54 controls (32). NHBE (**Fig. 1B**) and A549 (low MOI) (**Fig. 1C**) data were gathered from Supplementary Tables 2 and 1 (23). Genes with Status = Low or Outlier were not considered for analysis. All other datasets where analyzed from GEO data using DESeq2 by the UIowa Bioinformatics Core (**see Code Availability**). Genes with a p-value of zero were treated as having a p-value equal to the next lowest p-value in that dataset. Genes with p > 0.05 (-log(p) > 1.30) were considered statistically significant. Graphs were generated using GraphPad Prism v8.

### Cell culture

DBT, 17Cl-1, HEK293T, and HeLa cells expressing the MHV receptor carcinoembryonic antigen-related cell adhesion molecule 1 (a gift from Dr. Thomas Gallagher, Loyola University, Chicago, IL) were grown in Dulbecco’s modified Eagle medium (DMEM) supplemented with 10% fetal bovine serum (FBS), HEPES, sodium pyruvate, non-essential amino acids, L-glutamine, penicillin and streptomycin. To create BMDMs, bone-marrow cells were harvested from C57BL/6 mice and differentiated by incubating cells with 10% L929 cell supernatants and 10% FBS in Roswell Park Memorial Institute (RPMI) media for seven days. Cells were washed and replaced with fresh media every day after the 4^th^ day. For analysis of the NAD metabolome HEK293T cells were transfected with 1µg of pEGFP-C1 Empty Vector or pEGFP-C1-CMV-PARP10 using CalPhos Mammalian Transfection Kit (Takara Bio). 6 hours later the cells were treated with chemical treatments in DMEM + 10% FBS at 37°C 5% CO_2_ for 18h.

### Mice

Animal studies were approved by the University of Kansas Institutional Animal Care and Use Committee (IACUC) as directed by the Guide for the Care and Use of Laboratory Animals (Protocol #252-01). Anesthesia or euthanasia were accomplished using ketamine/xylazine. Pathogen-free C57BL/6 mice were purchased from Jackson Laboratories and maintained in the animal care facility at the University of Kansas.

### Virus infection

Recombinant WT (rJIA - GFP*rev*N1347) and N1347A (rJ-IA-GFP-N1347A) MHV were previously described (3). Both viruses expressed eGFP. MHV-A59 was previously described (44). All viruses were propagated on 17Cl-1 as previously described (11). DBT, 17Cl-1, and BMDM cells were infected as described in the figure legends with a 1-hour adsorption period, before virus was removed from the well and replaced with fresh media. For NAD analysis, DBT cells were washed with PBS and replenished with serum-free DMEM prior to infection with no supplements except penicillin and streptomycin and were maintained in serum-free media throughout the infection. For treatments with NAD modulating compounds, 10 μM NA, NAM, and SBI-797812 were added immediately following the adsorption phase. For NR experiments, 100 nmol of NR was added to cells in 24 well plates in serum-free media 4 hr prior to infection, removed during the adsorption phase, then another 100 nmol NR in serum free media was added following the adsorption phase. Another 100 nmol of NR was then added directly to the media again at 12 hpi. Cells and supernatants were collected at 18 hpi and viral titers were determined by plaque assay.

### Quantitative NAD metabolomics

NAD metabolites were quantified against internal standards in two LC-MS/MS runs as described (33).

### Western Analysis of PARP MARylation

HEK293T were transfected with 3µg pEGFP-C1 empty vector, pEGFP-C1-CMV-PARP7, pEGFP-C1-CMV-PARP10, pEGFP-C1-CMV-PARP12, or pEF1-EGFP-C1-PARP14[553-1801] via CalPhos Mammalian Transfection Kit. Six hours later the cells were treated with chemical treatments in DMEM + 10% FBS at 37°C 5% CO_2_ for 18h. Cells were washed in PBS and lysed in 50 mM HEPES pH 7.4, 150 mM NaCl, 1mM MgCl_2_, 1 mM TCEP, 1% Triton X-100 with the addition of Protease Inhibitors (Roche), 30 µM rucaparib (Selleck), and 1 µM PDD0017273 (Sigma). Lysates were microcentrifuged for 15 min at 4°C, quantified by Bradford assay, and supernatants were transferred to new tube with 4x SDS sample loading buffer (0.2M Tris-HCl pH 6.5, 4% BME, 8% w/v SDS, 0.0.8% Bromophenol Blue, 40% Glycerol). Samples were resolved via SDS-PAGE and transferred to nitrocellulose. Blots were blocked with 5% Milk-PBST for 30 min, incubated O/N in primary antibody (Rabbit Pan-ADPr 1:1000, Cell Signaling E6F6A; Rabbit GFP 1:1000, Chromotek PABG1-100; Mouse Tubulin 1:1000; Cell Signaling DM1A). Primary incubation was followed with HRP-conjugated secondary antibodies (Rabbit-HRP 1:10000, Jackson Laboratories 111-035-144; Mouse-HRP 1:5000, Invitrogen 62-6520). Blots were developed by chemiluminescence and imaged on a ChemiDoc MP system (Bio-Rad). Blot analysis was performed in Image-Lab (Bio-Rad).

### PARP Plasmids

pEGFP-C1-CMV-PARP7, pEGFP-C1-CMV-PARP10, pEGFP-C1-CMV-PARP12, and pEF1-EGFP-C1-PARP14[553-1801] were generated with standard restriction digest cloning.

### Data and code availability

Transcriptomic data for all NAD related genes are provided in Supplementary Materials 2-11. Primary RNAseq data are from publis (23,25)(32). NAD metabolomics data are provided in Supplementary Materials 13-14. Code for the generation of data in Supplementary Materials 2, 5-7, and 9-11 are provided.

## SUPPLEMENTARY MATERIALS

1. NAD gene list
2. Differential gene expression for Figure 1A
3. Differential gene expression for Figure 1B
4. Differential gene expression for Figure 1C
5. Differential gene expression for Figure 1D
6. Differential gene expression for Figure 1E
7. Differential gene expression for Figure 1F
8. Differential gene expression for Figure 2A
9. Differential gene expression for Figure 2B
10. Differential gene expression for Figure 2C
11. Differential gene expression for Figure 2D
12. NAD synthesis gene expression summarized Supplemental Materials 2-11.
13. NAD metabolomics for Figure 3A
14. NAD metabolomics for Figure 4A
15. NAD metabolomics for Figure 4B

## AUTHOR CONTRIBUTIONS

ARF, MSC and CB designed the experiments. YMOA, LSV, and DJS performed experiments with ARF and MSC, respectively. SAJT and MSS obtained NAD metabolomic data with CB. SP provided expertise. CDH performed informatic analyses with CB. Data were analyzed by all authors. The manuscript was written by CB with assistance of ARF, MSC and CDH.

## ACKNOWLEDGMENTS

We thank Stephen Gardell for kind gift of SBI-797812, ChromaDex for Niagen, Michael Chimenti and Henry Keen for informatic support, and Noah Fluharty and Grant Welk for assistance in graphing and statistics. Work was supported by NIH grants HL147545 (CB), GM008629 (SAJT), GM113117 and AI134993 (ARF), AI060699 and AI091322 (SP), CA245722 (CDH), 1NS08862 (MSC), Roy J. Carver Trust (CB), Alfred E. Mann Family Foundation (CB) and Pew Charitable Trusts (MSC).

## REFERENCES

1. Dong, E., Du, H., and Gardner, L.a (2020) An interactive web-based dashboard to track COVID-19 in real time. Lancet Infect Dis 20, 533–534

2. Wu, D., Wu, T., Liu, Q., and Yang, Z. (2020) The SARS-CoV-2 outbreak: what we know. International journal of infectious diseases : IJID : official publication of the International Society for Infectious Diseases 94, 44–48

3. Fehr, A. R., and Perlman, S. (2015) Coronaviruses: an overview of their replication and pathogenesis. Methods Mol Biol 1282, 1–23

4. Zhu, N., Zhang, D., Wang, W., Li, X., Yang, B., Song, J., Zhao, X., Huang, B., Shi, W., Lu, R., Niu, P., Zhan, F., Ma, X., Wang, D., Xu, W., Wu, G., Gao, G. F., Tan, W., China Novel Coronavirus, I., and Research, T. (2020) A Novel Coronavirus from Patients with Pneumonia in China, 2019. N Engl J Med 382, 727–733

5. Wu, C., Liu, Y., Yang, Y., Zhang, P., Zhong, W., Wang, Y., Wang, Q., Xu, Y., Li, M., Li, X., Zheng, M., Chen, L., and Li, H. (2020) Analysis of therapeutic targets for SARS-CoV-2 and discovery of potential drugs by computational methods. Acta Pharm Sinica B 10, 766–788

6. Belenky, P., Bogan, K. L., and Brenner, C. (2007) NAD+ metabolism in health and disease. Trends in Biochemical Sciences 32, 12–19

7. Alhammad, Y. M. O., and Fehr, A. R. (2020) The Viral Macrodomain Counters Host Antiviral ADP-Ribosylation. Viruses 12, 384

8. Fehr, A. R., Singh, S. A., Kerr, C. M., Mukai, S., Higashi, H., and Aikawa, M. (2020) The impact of PARPs and ADP-ribosylation on inflammation and host-pathogen interactions. Genes Dev 34, 341–359

9. Cohen, M. S. (2020) Interplay between compartmentalized NAD(+) synthesis and consumption: a focus on the PARP family. Genes Dev 34, 254–262

10. Gupte, R., Liu, Z., and Kraus, W. L. (2017) PARPs and ADP-ribosylation: recent advances linking molecular functions to biological outcomes. Genes Dev 31, 101–126

11. Grunewald, M. E., Chen, Y., Kuny, C., Maejima, T., Lease, R., Ferraris, D., Aikawa, M., Sullivan, C. S., Perlman, S., and Fehr, A. R. (2019) The coronavirus macrodomain is required to prevent PARP-mediated inhibition of virus replication and enhancement of IFN expression. PLoS Pathog 15, e1007756

12. Grunewald, M. E., Shaban, M. G., Mackin, S. R., Fehr, A. R., and Perlman, S. (2020) Murine Coronavirus Infection Activates the Aryl Hydrocarbon Receptor in an Indoleamine 2,3-Dioxygenase-Independent Manner, Contributing to Cytokine Modulation and Proviral TCDD-Inducible-PARP Expression. Journal of virology 94, e01743–01719

13. Fehr, A. R., Channappanavar, R., Jankevicius, G., Fett, C., Zhao, J., Athmer, J., Meyerholz, D. K., Ahel, I., and Perlman, S. (2016) The Conserved Coronavirus Macrodomain Promotes Virulence and Suppresses the Innate Immune Response during Severe Acute Respiratory Syndrome Coronavirus Infection. mBio 7, e01721–01716

14. Fehr, A. R., Athmer, J., Channappanavar, R., Phillips, J. M., Meyerholz, D. K., and Perlman, S. (2015) The nsp3 macrodomain promotes virulence in mice with coronavirus-induced encephalitis. Journal of virology 89, 1523–1536

15. Eriksson, K. K., Cervantes-Barragan, L., Ludewig, B., and Thiel, V. (2008) Mouse hepatitis virus liver pathology is dependent on ADP-ribose-1’’-phosphatase, a viral function conserved in the alpha-like supergroup. J Virol 82, 12325–12334

16. Li, L., Zhao, H., Liu, P., Li, C., Quanquin, N., Ji, X., Sun, N., Du, P., Qin, C. F., Lu, N., and Cheng, G. (2018) PARP12 suppresses Zika virus infection through PARP-dependent degradation of NS1 and NS3 viral proteins. Sci Signal 11, eaas9332

17. Atasheva, S., Akhrymuk, M., Frolova, E. I., and Frolov, I. (2012) New PARP gene with an anti-alphavirus function. J Virol 86, 8147–8160

18. Atasheva, S., Frolova, E. I., and Frolov, I. (2014) Interferon-stimulated poly(ADP-Ribose) polymerases are potent inhibitors of cellular translation and virus replication. J Virol 88, 2116–2130

19. Gardell, S. J., Hopf, M., Khan, A., Dispagna, M., Hampton Sessions, E., Falter, R., Kapoor, N., Brooks, J., Culver, J., Petucci, C., Ma, C. T., Cohen, S. E., Tanaka, J., Burgos, E. S., Hirschi, J. S., Smith, S. R., Sergienko, E., and Pinkerton, A. B. (2019) Boosting NAD(+) with a small molecule that activates NAMPT. Nat Commun 10, 3241

20. Trammell, S. A., Schmidt, M. S., Weidemann, B. J., Redpath, P., Jaksch, F., Dellinger, R. W., Li, Z., Abel, E. D., Migaud, M. E., and Brenner, C. (2016) Nicotinamide riboside is uniquely and orally bioavailable in mice and humans. Nat Commun 7, 12948

21. Dollerup, O. L., Christensen, B., Svart, M., Schmidt, M. S., Sulek, K., Ringgaard, S., Stodkilde-Jorgensen, H., Moller, N., Brenner, C., Treebak, J. T., and Jessen, N. (2018) A randomized placebo-controlled clinical trial of nicotinamide riboside in obese men: safety, insulin-sensitivity, and lipid-mobilizing effects. The American journal of clinical nutrition 108, 343–353

22. Elhassan, Y. S., Kluckova, K., Fletcher, R. S., Schmidt, M. S., Garten, A., Doig, C. L., Cartwright, D. M., Oakey, L., Burley, C. V., Jenkinson, N., Wilson, M., Lucas, S. J. E., Akerman, I., Seabright, A., Lai, Y. C., Tennant, D. A., Nightingale, P., Wallis, G. A., Manolopoulos, K. N., Brenner, C., Philp, A., and Lavery, G. G. (2019) Nicotinamide Riboside Augments the Aged Human Skeletal Muscle NAD(+) Metabolome and Induces Transcriptomic and Anti-inflammatory Signatures. Cell Rep 28, 1717–1728 e1716

23. Blanco-Melo, D., Nilsson-Payant, B. E., Liu, W.-C., Moller, R., Panis, M., Sachs, D., Albrecht, R. A., and tenOever, B. R. (2020) SARS-CoV-2 launches a unique transcriptional signature from in vitro, ex vivo, and in vivo systems. Cell 181, 1036–1045

24. Hoffmann, M., Kleine-Weber, H., Schroeder, S., Kruger, N., Herrler, T., Erichsen, S., Schiergens, T. S., Herrler, G., Wu, N. H., Nitsche, A., Muller, M. A., Drosten, C., and Pohlmann, S. (2020) SARS-CoV-2 Cell Entry Depends on ACE2 and TMPRSS2 and Is Blocked by a Clinically Proven Protease Inhibitor. Cell 181, 271–280 e278

25. Lamers, M. M., Beumer, J., van der Vaart, J., Knoops, K., Puschhof, J., Breugem, T. I., Ravelli, R. B. G., Paul van Schayck, J., Mykytyn, A. Z., Duimel, H. Q., van Donselaar, E., Riesebosch, S., Kuijpers, H. J. H., Schippers, D., van de Wetering, W. J., de Graaf, M., Koopmans, M., Cuppen, E., Peters, P. J., Haagmans, B. L., and Clevers, H. (2020) SARS-CoV-2 productively infects human gut enterocytes. Science 369, 50–54

26. Shi, J., Wen, Z., Zhong, G., Yang, H., Wang, C., Huang, B., Liu, R., He, X., Shuai, L., Sun, Z., Zhao, Y., Liu, P., Liang, L., Cui, P., Wang, J., Zhang, X., Guan, Y., Tan, W., Wu, G., Chen, H., and Bu, Z. (2020) Susceptibility of ferrets, cats, dogs, and other domesticated animals to SARS-coronavirus 2. Science 368, 1016–1020

27. Bieganowski, P., and Brenner, C. (2004) Discoveries of nicotinamide riboside as a nutrient and conserved NRK genes establish a preiss-handler independent route to NAD+ in fungi and humans. Cell 117, 495–502

28. Vaur, P., Brugg, B., Mericskay, M., Li, Z., Schmidt, M. S., Vivien, D., Orset, C., Jacotot, E., Brenner, C., and Duplus, E. (2017) Nicotinamide riboside, a form of vitamin B3, protects against excitotoxicity-induced axonal degeneration. FASEB J 31, 5440–5452

29. Diguet, N., Trammell, S. A. J., Tannous, C., Deloux, R., Piquereau, J., Mougenot, N., Gouge, A., Gressette, M., Manoury, B., Blanc, J., Breton, M., Decaux, J. F., Lavery, G., Baczko, I., Zoll, J., Garnier, A., Li, Z., Brenner, C., and Mericskay, M. (2018) Nicotinamide Riboside Preserves Cardiac Function in a Mouse Model of Dilated Cardiomyopathy. Circulation 137, 2256–2273

30. Neelakantan, H., Brightwell, C. R., Graber, T. G., Maroto, R., Wang, H. L., McHardy, S. F., Papaconstantinou, J., Fry, C. S., and Watowich, S. J. (2019) Small molecule nicotinamide N-methyltransferase inhibitor activates senescent muscle stem cells and improves regenerative capacity of aged skeletal muscle. Biochem Pharmacol 163, 481–492

31. Brenner, C. (2014) Metabolism: Targeting a fat-accumulation gene. Nature 508, 194–195

32. Lieberman, N. A. P., Peddu, V., Xie, H., Shrestha, L., Huang, M. L., Mears, M. C., Cajimat, M. N., Bente, D. A., Shi, P. Y., Bovier, F., Roychoudhury, P., Jerome, K. R., Moscona, A., Porotto, M., and Greninger, A. L. (2020) In vivo antiviral host transcriptional response to SARS-CoV-2 by viral load, sex, and age. PLoS Biol 18, e3000849

33. Trammell, S. A., and Brenner, C. (2013) Targeted, LCMS-based Metabolomics for Quantitative Measurement of NAD(+) Metabolites. Comput Struct Biotechnol J 4, e201301012

34. Yang, J., Zheng, Y., Gou, X., Pu, K., Chen, Z., Guo, Q., Ji, R., Wang, H., Wang, Y., and Zhou, Y. (2020) Prevalence of comorbidities in the novel Wuhan coronavirus (COVID-19) infection: a systematic review and meta-analysis. International journal of infectious diseases : IJID : official publication of the International Society for Infectious Diseases 94, 91–95

35. Trammell, S. A., Weidemann, B. J., Chadda, A., Yorek, M. S., Holmes, A., Coppey, L. J., Obrosov, A., Kardon, R. H., Yorek, M. A., and Brenner, C. (2016) Nicotinamide Riboside Opposes Type 2 Diabetes and Neuropathy in Mice. Scientific reports 6, 26933

36. Kunzi, L., and Holt, G. E. (2019) Cigarette smoke activates the parthanatos pathway of cell death in human bronchial epithelial cells. Cell Death Discov 5, 127

37. Liu, H. W., Smith, C. B., Schmidt, M. S., Cambronne, X. A., Cohen, M. S., Migaud, M. E., Brenner, C., and Goodman, R. H. (2018) Pharmacological bypass of NAD(+) salvage pathway protects neurons from chemotherapy-induced degeneration. Proc Natl Acad Sci U S A 115, 10654–10659

38. Murray, M. F., Nghiem, M., and Srinivasan, A. (1995) HIV infection decreases intracellular nicotinamide adenine dinucleotide [NAD]. Biochem Biophys Res Commun 212, 126–131

39. Grady, S. L., Hwang, J., Vastag, L., Rabinowitz, J. D., and Shenk, T. (2012) Herpes simplex virus 1 infection activates poly(ADP-ribose) polymerase and triggers the degradation of poly(ADP-ribose) glycohydrolase. J Virol 86, 8259–8268

40. Bogan, K. L., and Brenner, C. (2008) Nicotinic acid, nicotinamide, and nicotinamide riboside: A molecular evaluation of NAD + precursor vitamins in human nutrition. Annual Review of Nutrition 28, 115–130

41. Rankin, P. W., Jacobson, E. L., Benjamin, R. C., Moss, J., and Jacobson, M. K. (1989) Quantitative studies of inhibitors of ADP-ribosylation in vitro and in vivo. J Biol Chem 264, 4312–4317

42. Chen, J., Sysol, J. R., Singla, S., Zhao, S., Yamamura, A., Valdez-Jasso, D., Abbasi, T., Shioura, K. M., Sahni, S., Reddy, V., Sridhar, A., Gao, H., Torres, J., Camp, S. M., Tang, H., Ye, S. Q., Comhair, S., Dweik, R., Hassoun, P., Yuan, J. X., Garcia, J. G. N., and Machado, R. F. (2017) Nicotinamide Phosphoribosyltransferase Promotes Pulmonary Vascular Remodeling and Is a Therapeutic Target in Pulmonary Arterial Hypertension. Circulation 135, 1532–1546

43. Mehta, P., McAuley, D. F., Brown, M., Sanchez, E., Tattersall, R. S., Manson, J. J., and Hlh Across Speciality Collaboration, U. K. (2020) COVID-19: consider cytokine storm syndromes and immunosuppression. Lancet 395, 1033–1034

44. Yount, B., Denison, M. R., Weiss, S. R., and Baric, R. S. (2002) Systematic assembly of a full-length infectious cDNA of mouse hepatitis virus strain A59. Journal of virology 76, 11065–11078

